# Reciprocal inflammatory signaling in an ex-vivo explant model for neurofibromatosis type 1-related cutaneous neurofibromas

**DOI:** 10.1101/2022.10.06.510849

**Authors:** Jamie Grit, Lisa Turner, Curt Essenburg, Patrick Dischinger, Nate Shurlow, Matt Pate, Carrie Graveel, Matt Steensma

## Abstract

Cutaneous neurofibromas (CNF) are benign tumors that occur in the dermis of individuals with neurofibromatosis type 1 (NF), an inherited tumor predisposition syndrome. CNFs cause disfigurement, pain, burning, and itching, resulting in reduced quality of life in NF patients. However, due to their benign nature there are few *in vitro* or *in vivo* models of CNFs, which has limited the research of CNF biology and drug discovery efforts. To address this, we developed a patient derived explant (PDE) *ex vivo* culture model of CNF tumors and normal skin from NF patients. CNF PDEs remain viable in culture for over 9 days and recapitulate the cellular composition and molecular signaling of CNFs. We identified reciprocal inflammatory signaling in CNF PDEs, in which tumors rely on either prostaglandin or leukotriene mediated signaling pathways. *Ex vivo* glucocorticoid treatment reduced expression of pro-inflammatory genes, confirming CNF PDEs are a useful model for mechanistic studies and preclinical drug testing.

## Introduction

Cutaneous neurofibromas (CNF) are defining features of the inherited tumor predisposition disorder, Neurofibromatosis Type 1 (NF). CNFs account for substantial morbidity, negatively impacting the quality of life of NF patients. Depending on the degree of skin involvement, NF-related CNFs are associated with disfigurement, intractable pain, burning and itching. For NF patients with a high density of CNFs, there are no effective treatments resulting in life-long morbidity. Despite the recent success in the treatment for NF-related plexiform neurofibromas (PNF) using the oral MEK inhibitor, selumetinib, data suggest that CNFs are biologically distinct tumors with respect to MEK inhibitor therapy (Dombi, Baldwin et al. 2016, Brosseau, Pichard et al. 2018, Verma, Riccardi et al. 2018, Gross, Wolters et al. 2020). We recently described differential signaling paradigms between PNFs and CNFs as a result of diverse Ras signaling outputs (Grit, Johnson et al. 2021). The therapeutic relevance of these findings is that RAS signaling is epigenetically reinforced to drive inflammation in CNFs (e.g. MKK3-p38) moreso than growth and proliferation (e.g. RAS-ERK), an observation that may partially explain the lack of selumetinib effectiveness in treating CNFs.

Interestingly, the effectiveness of selumetinib in treating PNFs was described simultaneously in a genetically engineered mouse model (DhhCre;Nf1fl/fl) and in a phase 1 human clinical trial (Dombi, Baldwin et al. 2016). At that time, there were no “humanized” orthotopic models of PNFs, so demonstration of this landmark drug response required simultaneous “proof of principle” experiments in a mouse model and direct treatment of human subjects. Today, there remains a gap in orthotopic neurofibroma models which could be leveraged towards drug prediction. The current animal models of NF-related CNFs are robust for biological studies because they recapitulate key features of the distinctive CNF phenotype, but there are significant limitations in using these models to translate findings into the clinic. Genetically engineered models (GEMs) of CNFs are typically not scalable due expense, prolonged time frames for spontaneous tumor evolution, and general limitations in extrapolating murine CNF biology to human disease.

In order to bridge the gap in translating animal CNF findings to human disease and to reduce reliance on animal testing, we propose the use of *ex vivo,* human CNF explants for drug testing, mechanistic studies, and correlative science from CNF-specific clinical trials which are currently ongoing. We present methodology and in-depth characterization of CNF explants demonstrating maintenance of tumor and cellular phenotype within ideal experimentation time frames (>9 days). Moreover, we employ our explant model to extend our prior findings regarding differences between downstream RAS signaling in CNFs versus PNFs. We further demonstrate that epigenetically reinforced CNF inflammatory signaling is both targetable and dichotomous, driven by expression changes in key mediators of prostaglandin or leukotriene synthesis.

## Results

### CNF PDEs are viable in culture for 9 days

CNFs are relatively small skin tumors comprised of spindle cells within a collagenous stroma. The major cellular components are S100B positive neoplastic neural lineage cells and CD34 positive dermal fibroblasts (Figure 1a). CNFs are contained within the dermis layer of the skin, thus we reasoned they could be cultured *ex vivo* similar to previously established skin explant culture models (Corzo-León, Munro et al. 2019, Powley, Patel et al. 2020, Rakita, Nikolić et al. 2020). CNF tumors (Figure 1a & b) were surgically excised (Figure 1c) and transferred to culture conditions (Figure 1d) within 15 minutes. Medium was carefully added so that the tumor/dermis layers were submerged and the epidermis remained dry at the air-liquid interface (Figure 1e), mimicking normal skin conditions.

**Figure 1.**
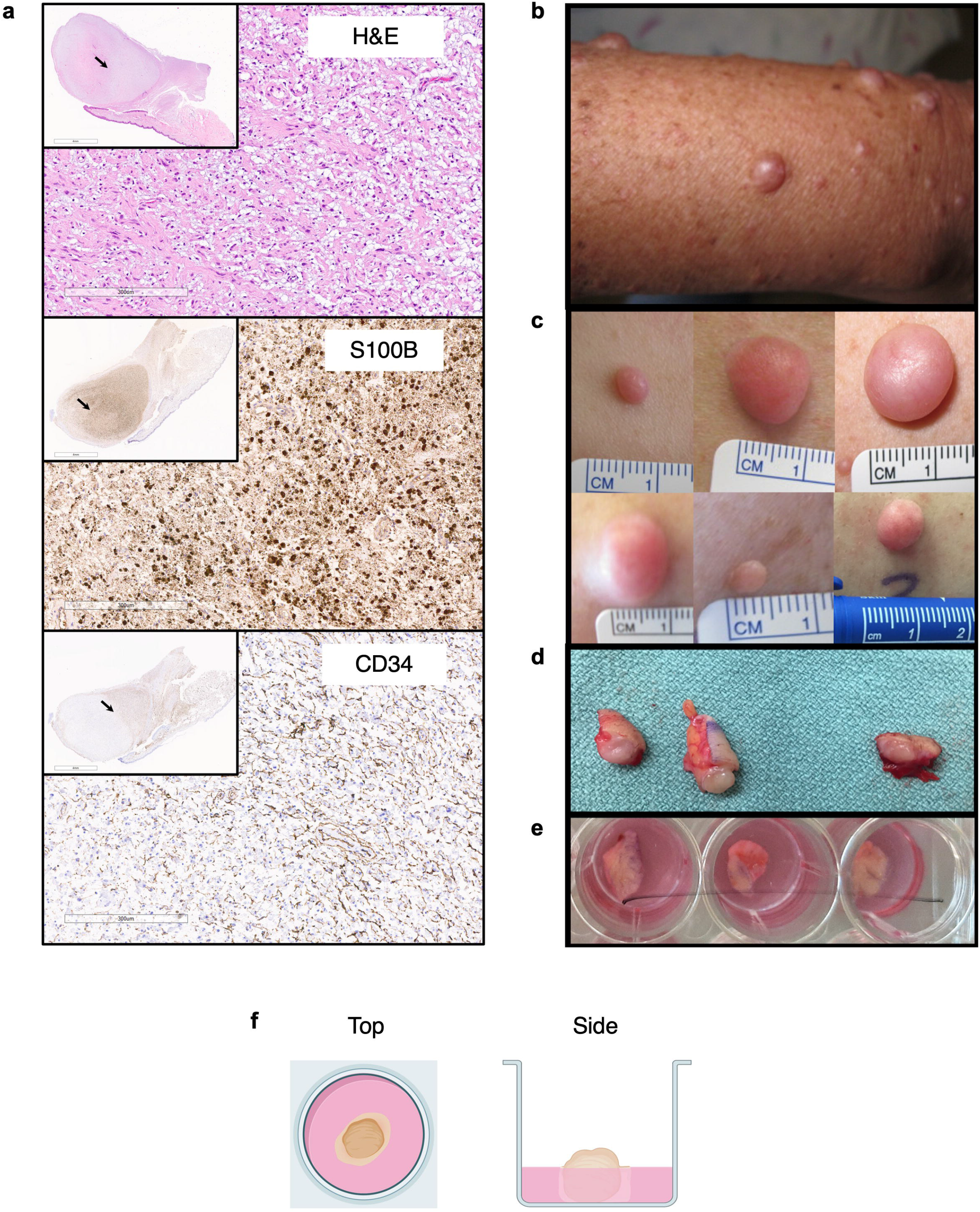
Generation of CNF PDEs with phenotype characterization. (A) H/E, S100B (neural lineage marker) and CD34 (fibroblast marker) immunohistochemistry staining shown. (B) Representative clinical presentation of multiple CNFs in the proximal forearm. (C) Clinical photographs of CNFs with size variation. (D) CNF tumor specimens immediate post resection. (E) CNF PDEs in 12 well tissue culture dish (3 wells shown). (F) Schematic of explant culture position in top down and side views.

Previous studies demonstrated skin explants can be maintained in culture for up to two weeks with minimal cell death (Corzo-León, Munro et al. 2019). We evaluated tissue integrity and cell viability in CNF PDEs by comparing histology and cleaved caspase 3 (CC3) staining for apoptotic cells at baseline and after extended time in culture. H&E staining revealed tumors comprised primarily of spindle cells (Figure 2a, top insets) within a collagenous stroma (Figure 2a, bottom insets) at each time point. There was no evidence of necrosis in PDEs even after 7 days in culture (n=12) with some tumors being cultured for up to 11 days successfully. CC3 staining demonstrated minimal apoptosis in cultured CNFs, even after one week or more in culture (n=12) (Figure 2b). CC3 positive cells were nearly exclusively keratinocytes within the stratum basale layer of the epidermis (Figure 2b, top insets) with very few apoptotic cells within the CNF itself (Figure 2b, bottom insets). CC3 staining in basale keratinocytes was similar across CNF PDEs regardless of time in culture. We next compared CC3 staining in CNF PDEs with CNFs that were formalin fixed within 30 minutes of surgical resection. Interestingly, CC3 positive keratinocytes were also present in the stratum basale layer of freshly fixed CNFs, although to a lesser degree (Supplemental Figure 1). This is consistent with findings by Simon et al, who showed CC3 positive keratinocytes are also found in the basal layer of patients with atopic dermatitis (Simon, Lindberg et al. 2006). Collectively, these data indicate that CNFs can be cultured *ex vivo* without significant changes in skin composition or viability.

**Figure 2.**
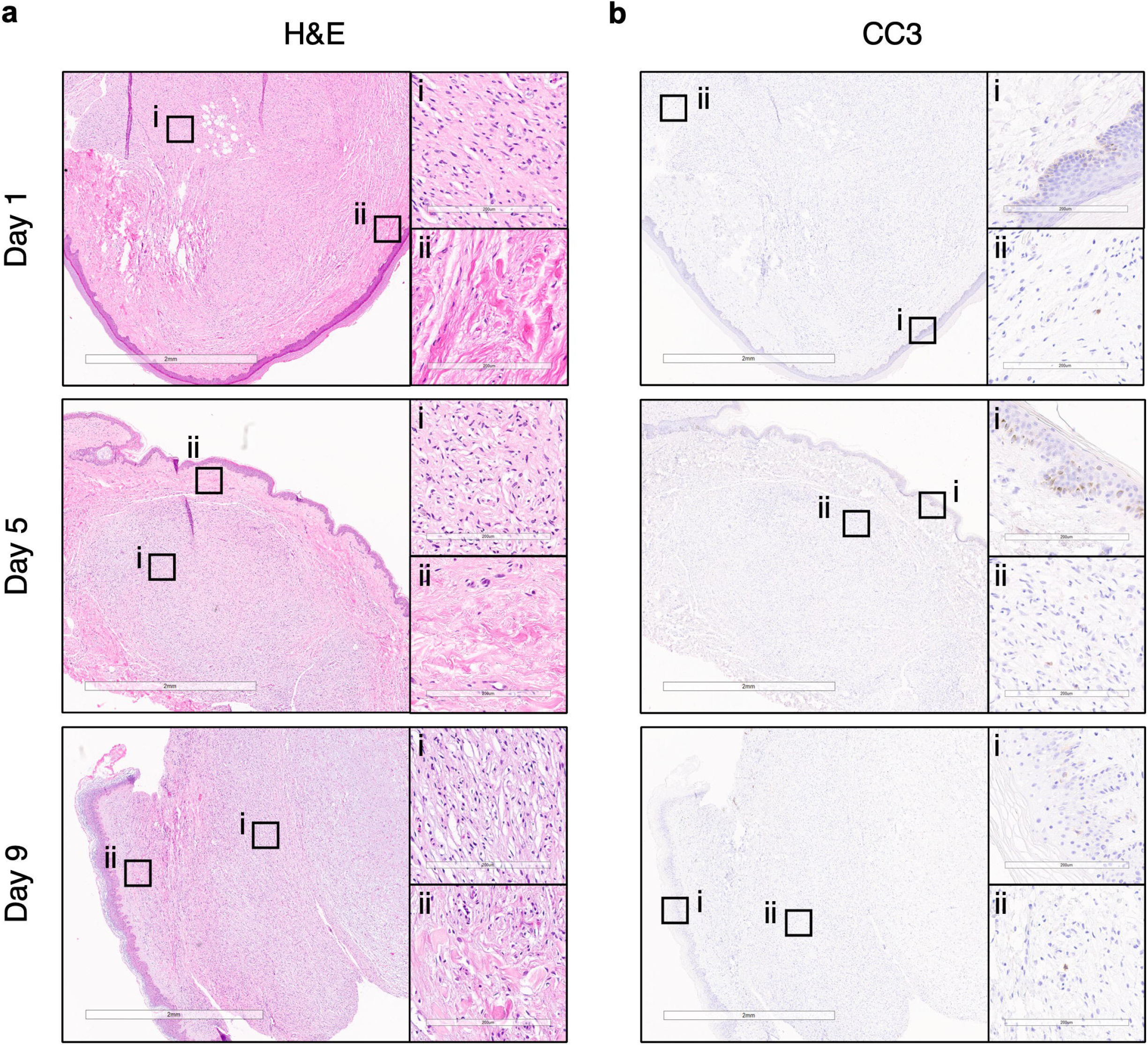
CNF PDEs are viable and retain structural architecture. (a) H&E staining (20x) of CNF PDEs after one, five, and nine days in culture. Top insets (i): characteristic spindle cells (200x); Bottom insets (ii): collagenous stroma (200x). (b) Cleaved caspase 3 IHC (20x) of CNF PDEs over time. Top insets (i): epidermis (200x); Bottom insets (ii): tumor region (200x).

CNFs are comprised of neoplastic, neural lineage cells as well as fibroblasts, mast cells, and macrophages, thus an ideal CNF model should recapitulate the complex tumor microenvironment. We evaluated CNF PDEs by IHC for the presence of neural lineage cells (S100B), dermal fibroblasts (CD34), mast cells (c-KIT), and macrophages (IBA1). As expected, S100B positive tumor cells and CD34 positive dermal fibroblasts were the most abundant cell types in the PDEs, regardless of time spent in culture (Figure 3a & b). Tumor cell and dermal fibroblast density in the CNF PDEs were highly variable among and within tumors, with regions of high and low S100B and CD34 expression both within individual tumors (Supplemental Figure 2) and across tumors, even within the same patient (Figure 3a & b). Both intra- and inter-tumoral immune cell infiltrate was also highly variable, with some tumors/regions showing dense infiltration of c-KIT positive mast cells and/or IBA1 positive macrophages, while staining was sparse or absent in others (Figure 3c & d). T- and B-cell populations were not directly assessed. These data support the use of PDEs as a model for CNF research that requires a complex tumor microenvironment. They also highlight the need for multiple tumor replicates, particularly in the study of inflammation and immunology.

**Figure 3.**
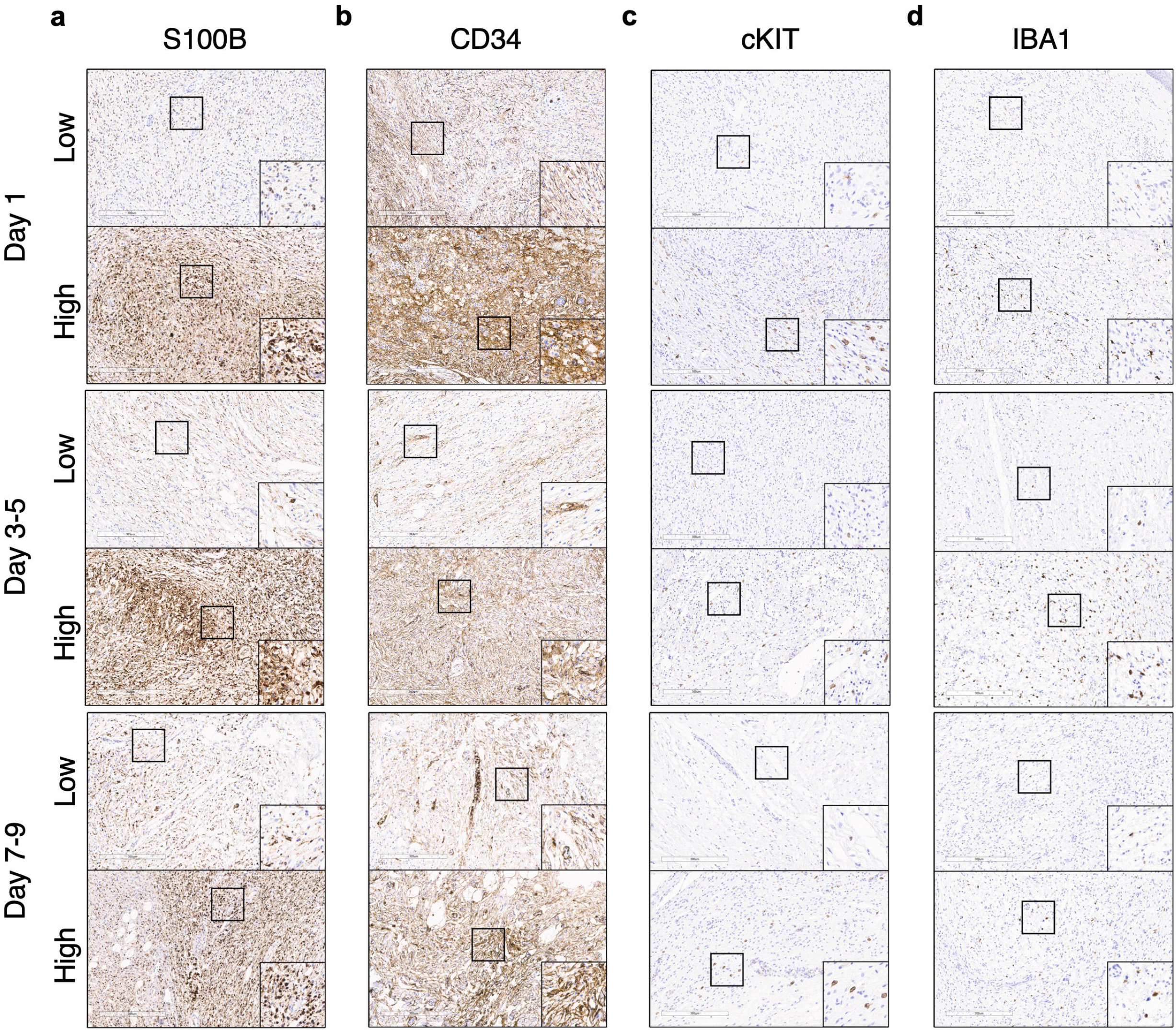
CNF PDEs model cellular heterogeneity. S100B positive tumor cells (a), CD34 positive dermal fibroblasts (b), cKIT positive mast cells (c), and IBA1 positive macrophages (d) (100x) after one day, 3-5 days, or 7-9 days in culture. Top panels show representative images of low expressing tumors, lower panels show images of high expression tumors. Insets 200x.

We next evaluated PDEs generated from the unaffected skin of NF1 patients and compared them to CNF PDEs. H&E staining revealed that the histology of skin PDEs was similar to that of uncultured skin (Figure 4a). Like CNF PDEs, there were very few apoptotic cells in the skin PDEs. Even after one week in culture, CC3 positive cells were largely restricted to keratinocytes of the stratum basale layer and nearly absent within the dermis (Figure 4b). Unlike CNF PDEs, expression of S100B in skin PDEs was restricted to melanocytes, Langerhans cells, and Schwann cells, similar to uncultured skin (Figure 4c, top panels). CD34 positive dermal fibroblasts were present in skin PDEs (Figure 4c, bottom panels), with lighter, more uniform staining than was observed in CNF PDEs (Figure 3b). These data show that skin PDEs from NF1 patients can be maintained in culture *ex vivo,* and suggest their use as a model for understanding how the *NF1* haploinsufficent microenvironment contributes to CNF origin and progression.

**Figure 4.**
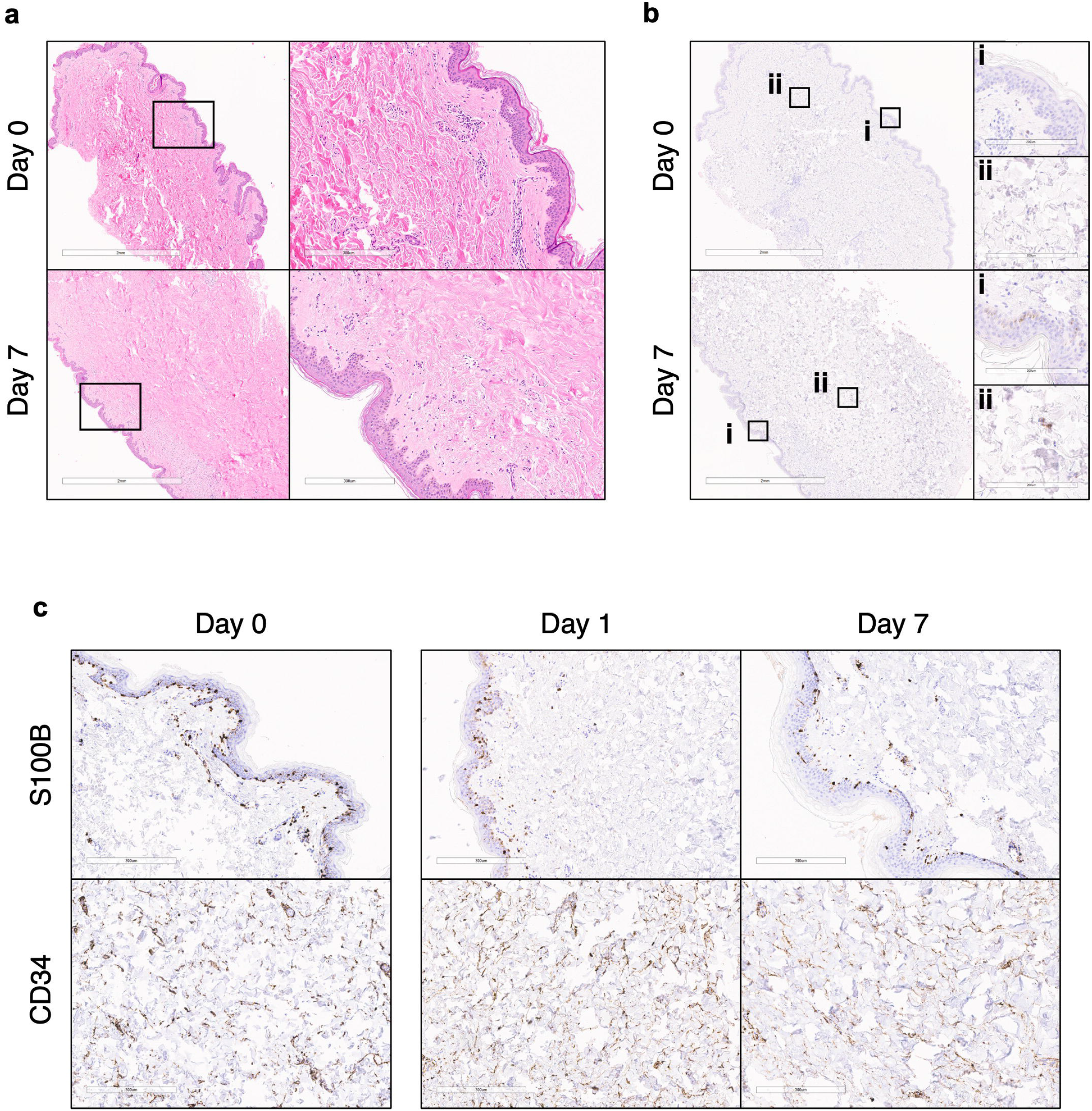
Normal skin PDEs from NF1 patients are viable in culture. (a) H&E staining (20x), insets (100x) of unaffected skin PDEs at baseline and after one week in culture. (b) Cleaved caspase 3 staining (20x) of unaffected skin PDEs at baseline and after one week in culture. Top insets (i) epidermis (200x); Bottom insets (ii) dermis (200x). (c) S100B (top) and CD34 (bottom) IHC of unaffected skin PDEs at baseline and after one and seven days in culture (100x).

We previously showed that CNFs have highly variable ERK activation, with phospho-ERK being absent in some CNFs and abundant in others (Grit, Johnson et al. 2021). Moreover, even within tumors, pERK staining is highly variable apart from consistent expression within the epidermis and vessel lining endothelial cell populations. We next wanted to determine whether CNF PDEs recapitulate this signaling heterogeneity, and if signaling heterogeneity is stable over time in culture. IHC for phospho-ERK revealed a wide range of expression across tumors, with some tumors negative for phospho-ERK (Figure 5, top panel) and others ranging from low to high expression (Figure 5 middle and lower panels). This heterogeneity was maintained in culture over time, indicating that CNF PDEs accurately model cell signaling events. Thus, CNF PDEs could conceivably be used for preclinical testing of kinase inhibitors, including topical formulations which could be applied to the epidermis, or systemic formulations which could be added to the culture medium.

**Figure 5.**
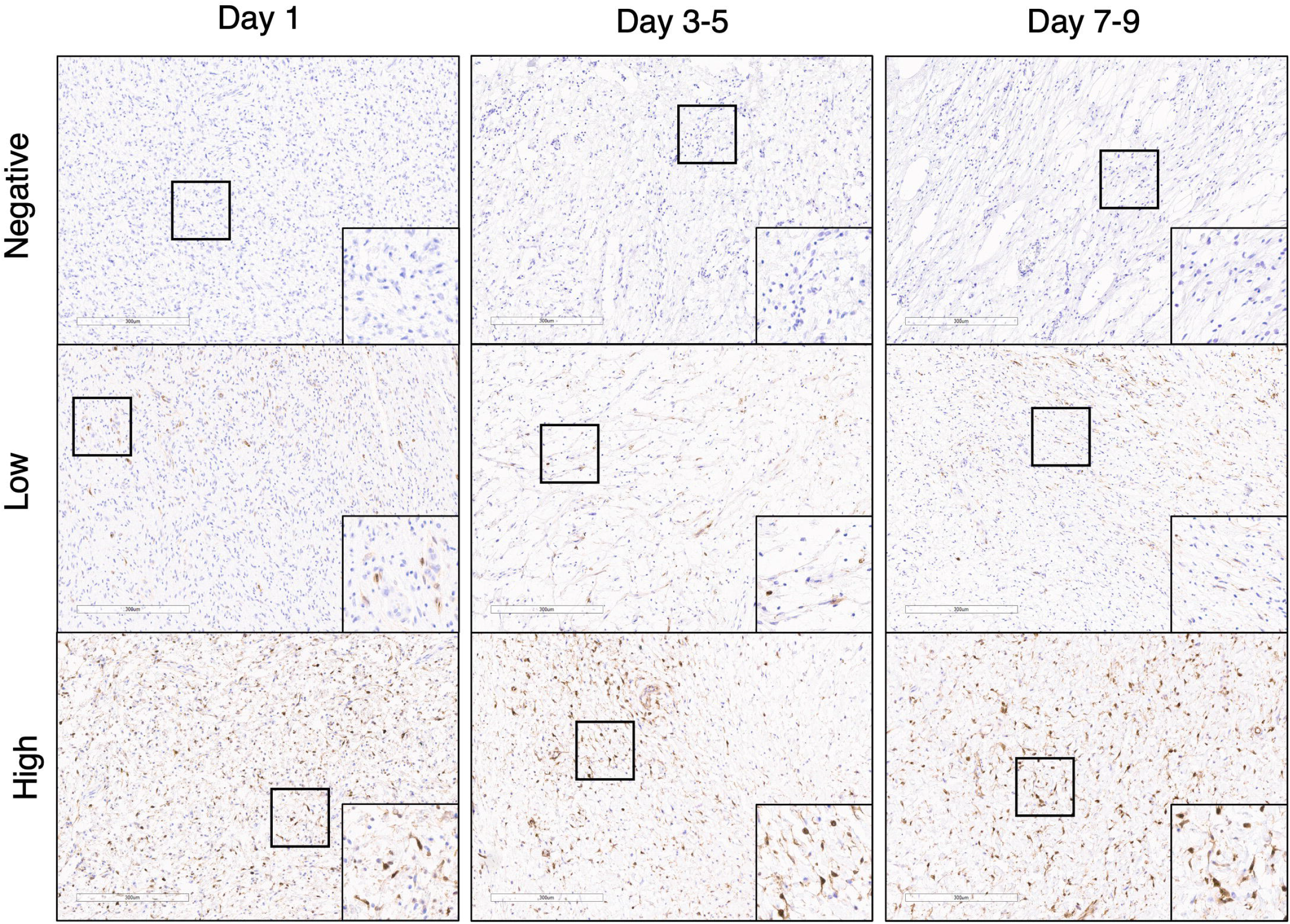
CNF PDEs model variable ERK activation. Phospho-ERK IHC (100x) after one day, 3-5 days, or 7-9 days in culture. Top panels show representative images of tumors negative for phospho-ERK expression, middle panels show tumors with low phospho-ERK expression, lower panels show tumors with high phospho-ERK expression. Insets 200x.

### Inflammation is decreased upon *ex vivo* drug treatment of CNF PDEs

We previously found increased inflammation in CNFs compared to PNFs (Grit, Johnson et al. 2021). The enzyme COX2 is an inflammatory mediator through the production of prostaglandin. COX2 activity is enhanced by iNOS mediated S-nitrosylation (Kim, Huri et al. 2005). We evaluated expression of both S-nitrosylated and non-nitrosylated COX2 by IHC (Jindal, Pennock et al. 2020) in CNF PDEs. COX2 expression was highly variable within and between patients, with both S-nitroslyated (Figure 6a) and non-nitrosylated (Figure 6b) forms of COX2 detected to varying degrees in tumor and adjacent tissue.

**Figure 6.**
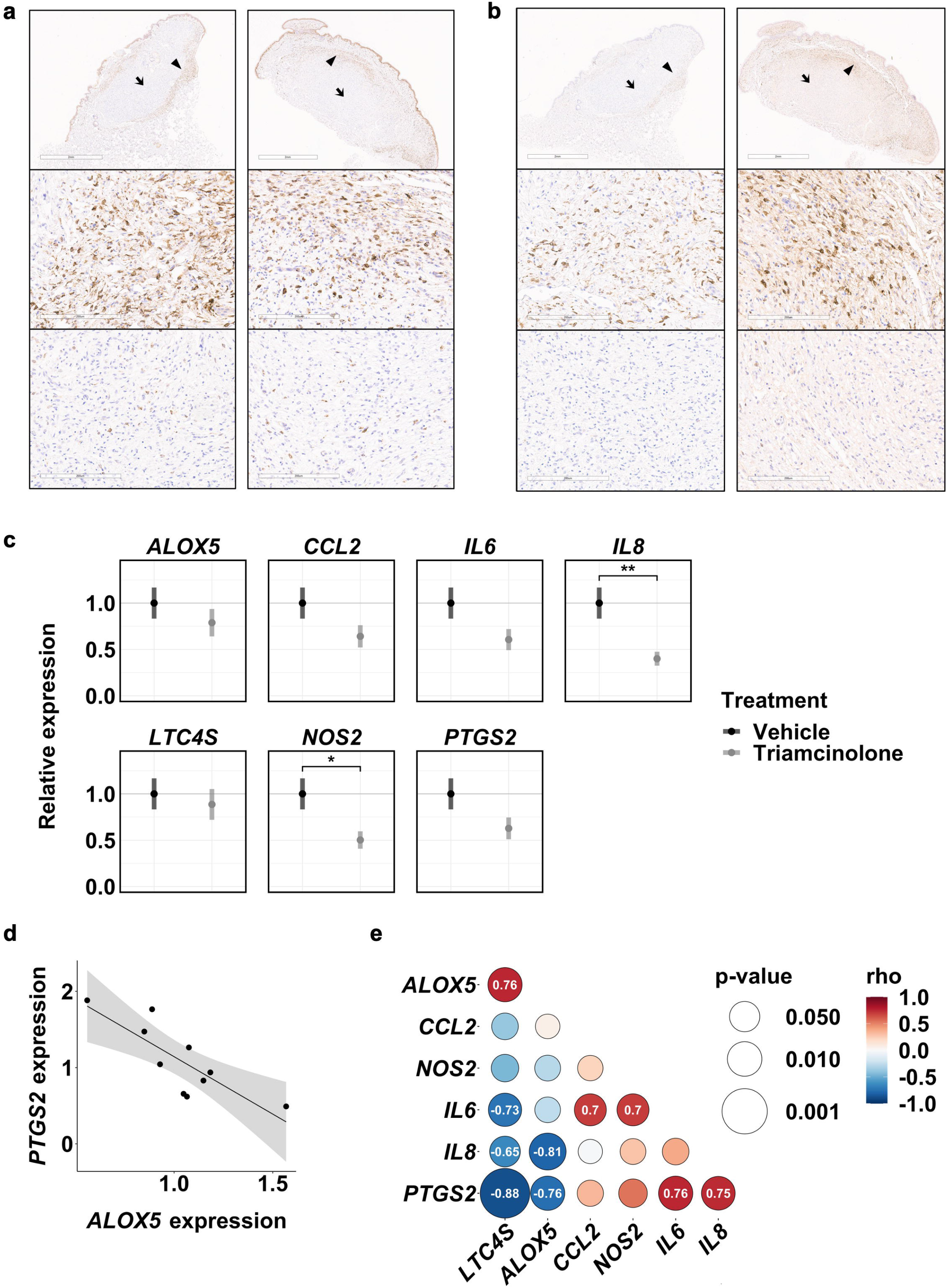
*Ex vivo* glucocorticoid injection reduces inflammation in CNF PDEs. (a) S-nitrosylated and (b) non-nitrosylated COX2 IHC (15x) in CNF PDEs. Arrowheads indicate regions of high expression (middle panels, 200X) and arrows indicate regions of low expression (lower panels, 200x). (c) Expression (qRT-PCR) of inflammatory markers in PDEs after glucocorticoid (triamcinolone) injection relative to vehicle. Points and error bars indicate the estimated marginal mean +/- SE. (d) Negative correlation between *PTGS2* and *ALOX5* expression in control PDEs; r = −0.81, p value = 0.004, shaded ribbon represents 95% confidence interval. (e) Correlogram depicting positive (red) and negative (blue) correlations in the expression of inflammatory markers in control PDEs. Dot color indicates Spearman’s rho, dot size indicates p-value. For significant correlations rho is indicated in white text within the dot.

We next used CNF PDEs as a preclinical model to test the anti-inflammatory drug triamcinolone. Triamcinalone is an injectable glucocorticoid approved for the treatment of several inflammatory skin conditions. Twenty-four PDEs were derived from CNFs harvested from a single patient and injected *ex vivo* with either triamcinolone or vehicle. After 24 hours gene expression of several inflammatory mediators, including *PTGS2* (COX2) and *NOS2* (iNOS) was evaluated. We found downregulation of all inflammatory markers, with *NOS2* and *IL8* being significantly decreased in triamcinolone treated PDEs (Figure 6c). These data show that CNF PDEs can be used to model gene expression changes in response to *ex vivo* stimuli, and also suggest that triamcinolone can reduce CNF inflammation.

Interestingly, in vehicle treated control PDEs, we observed an inverse correlation between *PTGS2* (COX2) and *ALOX5* (5-lipoxygenase) expression (Figure 6d). Arachidonic acid is the substrate for both COX2 (which generates prostaglandins) and 5-lipoxygenase (which generates leukotrienes). These data suggest that CNFs rely either on prostaglandin (COX2) driven inflammation or leukotriene (5-lipoxygenase) driven inflammation. We evaluated whether any other of the inflammatory markers we profiled were associated with *PTGS2* or *ALOX5* expression. Expression of *LTC4S,* the enzyme downstream of *ALOX5,* was strongly correlated with *ALOX5* expression, while cyto/chemokines (*IL6, IL8, CCL2*) and *NOS2* were elevated in high *PTGS2* expressing tumors (Figure 6e).

## Discussion

We demonstrate a tractable explant model of NF-related CNFs. Based on a comprehensive review of the literature, this model is the first description of a benign, skin tumor explant. The feasibility of normal skin explants is well established (Eberlin, Silva et al. 2020, Neil, Brown et al. 2020, Eberlin, Facchini et al. 2021). Skin explants have been useful for studying drug response, cellular biology and for post-hoc cell isolation. In these models, data confirms maintenance of both the epidermal, dermal and tumor phenotype in culture beyond 9 days. In our CNF explant model, we confirm both viability and maintenance of cellular phenotypes at 9 days (Figures 2, 3 & 5) which is a sufficient window for studying drug responses, cellular composition, and molecular signaling. Interestingly, cellular composition was maintained over the 9 day observation time frame despite observed inter-tumoral heterogeneity (Figure 3). The sustained presence of mast cells and tissue macrophages in tissue culture, however, suggest that the explants do maintain at least some immunologic function in culture.

Explant models are generally limited by the availability of human tissue, but surgical resection of NF-related CNFs is common, particularly in centers with active dermato-oncology or surgical oncology practices. By age 70, 100% of NF-affected individuals demonstrate symptomatic CNFs (Tabata, Li et al. 2020) and the severity of CNF-related symptoms often necessitates treatment. Thus, CNF PDEs can be derived in sufficient numbers from patients or research subjects in clinical trials. The lack of effective treatments for NF-related CNFs, particularly in patients with a high tumor burden, has led to a reinvigorated effort on developing new therapeutic approaches for NF-related CNFs (Verma, Riccardi et al. 2018). An *ex vivo,* human tumor model would be extremely useful for predicting therapy responses, to analyze on-target drug effects, to perform drug screening experiments on human tissue, for tumor and normal skin cell isolation, to assess drug toxicity, and to define targetable signaling paradigms.

As such, we examined the utility of our explant model to assess whether it accurately recapitulates human disease. First, we assessed downstream markers of RAS-MKK3-P38 versus RAS-ERK signaling in NF-related CNF explants (Grit, Johnson et al. 2021). We discovered elevated expression of key mediators of prostaglandin (*PTGS2*) and leukotriene synthesis (*ALOX5*) (Figure 6) among multiple CNF explants. Interestingly, an inverse correlation was noted between PTGS2-driven and ALOX5-driven explants suggesting that NF-related CNFs can be uniquely dependent on prostaglandins or leukotrienes, but typically not both. When we simultaneously assessed RAS-ERK activation (pERK), some CNF explants failed to express basal pERK expression apart from scant expression in endothelial cells, whereas others demonstrate abundant pERK expression across multiple cell types. The reasons for this signaling heterogeneity is unclear, but it does underscore the need for additional treatment options apart from MEK inhibitors for NF-related CNFs.

These data also confirm our prior work that deregulated RAS signaling in CNFs is epigenetically reinforced to drive the differential expression of genes involved in p38-mediated inflammation, such as arachidonic acid metabolism. The moderately potent corticosteroid, triamcinolone is known to directly inhibit p38, COX2 and 5-lipoxygenase (Zhang, Lai et al. 2013, Johnstone, Honeycutt et al. 2019). As expected, triamcinolone treatment of CNF explants inhibited or suppressed markers of downstream activation of the RAS-MKK3-p38 inflammatory signaling axis, most notably *IL-8* and *NOS2.* Based on IHC staining of the explant specimens, COX2 expression was variable but demonstrated in both the dermis and tumor. DNA was extracted from homogenized specimens meaning that both tumor and dermis DNA was analyzed simultaneously, so the inverse *IL-8NOS2* expression observation was robust despite sampling of mixed tissues. These data suggest that triamcinolone treatment can be used to alter inflammatory signaling within the tumor and adjacent skin. Given that the CNF cell of origin resides in the skin microenvironment, these findings are important to justifying broader testing of potent corticosteroid injections in skin tumors.

Given the success in inhibiting RAS-mediated inflammation in CNF explants with triamcinolone injection, topical steroid treatment also seems to be an enticing strategy (https://clinicaltrials.gov/ct2/show/NCT04435665). Although we didn’t conduct a topical corticosteroid experiment, there is now sufficient justification to do so. Topical drug administration is feasible with human skin explant models, as well as our CNF explant culturing technique (Figure 1) (Eberlin, Silva et al. 2020). When these types of experiments are performed with matching control skin from tissue donors, the explants are essentially isogenic with respect to NF1 gene status (Figure 4), making them ideal models to address both inter- and intratumoral heterogeneity.

## Conclusion

NF-related CNF explants remain viable for 9 days and are a tractable model for biology studies, drug screening, and correlative science. RAS-mediated inflammation is dependent on COX2 and ALOX5 which can all be inhibited effectively by triamcinolone. CNF signaling downstream of p38 is diverse with an inverse relationship between *PTGS2* and *ALOX5* expression. More work is needed to verify the therapeutic effects of inhibiting inflammation in NF-related CNFs, however the degree of variable pERK expression in CNF PDEs and human clinical samples suggests that CNFs will not respond similarly to PNFs when treated with MEK inhibitors.

## Methods

### Tissue collection

Research subjects were enrolled for tumor and tissue harvesting in accordance with standardized IRB protocols and practice (IRB#2011-01). Inclusion criteria were designed to evaluate NF-related CNFs of varying morphology and anatomic locations. Research subjects over the age of 18 with a confirmed diagnosis of NF1 and clinically identifiable CNFs were allowed to participate. CNFs with ulcerations, atypical features, or more deeply situated masses (e.g. diffuse neurofibromas involving skin) were excluded. CNF tumors were excised as part of routine care using standard sterile technique. Routine pathologic assessment by a board-certified pathologist was performed to verify diagnosis. The surgical technique was standardized for all specimens. Briefly, the skin was prepped with a 4% chlorhexidine solution. 0.25% Marcaine was injected at the base of each lesion taking care not to penetrate the tumor, itself. A #15 blade scalpel was used to excise a small margin of skin at the base of each lesion. Bovie electrocautery was withheld until after the mass was removed to avoid thermal necrosis of the specimen. A thin layer of subcutaneous tissue was also removed as deep margin. Once the specimen was isolated from the surgical field, it was placed in a sterile container on a saline-soaked non-adhesive pad and placed on wet ice for transport. Within 15 minutes, the specimen(s) were transported to the laboratory for culturing. Using sterile technique, each explant was carefully placed into culture medium.

### Culture

Tumors smaller than 5 mm were placed cut side down directly in 12 well tissue culture plates using aseptic technique in a biosafety cabinet. Larger tumors were dissected into smaller pieces using sterile surgical tools prior to culturing. Tumors with a very uneven cut surface were placed directly against the well wall to help them remain upright. DMEM with 10% FBS and 1% pen/strep was carefully added to the well so that the tumor/dermis layers were submerged and the epidermis remained dry at the air-liquid interface. Tumors were placed in the incubator at 5% CO2, and medium changed as needed (daily or every other day) for the duration of the experiment.

### Histology

Tissues were fixed in 10% neutral buffered formalin for 72 hours (Thermo) then paraffin embedded and sectioned for histology and immunohistochemistry. Deparaffinization and antigen retrieval was performed on Dako PT link platform using Dako High pH retriever buffer for 20 minutes at 97 degrees C. Staining was performed utilizing Dako Autostainer Link 48, utilizing Dako Rabbit Polymer HRP or Dako EnVision Flex Polymer for secondary for 20 minutes following primary antibody incubation for 30 minutes. Primary antibodies were as followers: CC3 (Asp175): Cell Signaling 9661 (1:100); S100B: Cell signaling 90393 (1:500); CD34: Cell Signaling 3569 (1:50); c-KIT: Cell Signaling 37805; IBA1: Cell Signaling 17198 (1:800), phospho-ERK (T202/Y204): Cell Signaling 4370 (1:800); nitrosylated-COX2: Invitrogen MA5014568 (1:100); total COX2: Cayman 160112 (1:100). DAB detection was performed using Dako EnVision Flex Chromagen for 10 minutes and Dako Flex Hematoxylin for 5 minutes. H&E staining was performed on the Tissue-Tek Prisma Plus Autostainer utilizing Tissue-Tek H&E Stain Kit #1. Aperio scanning was performed utilizing the Leica Aperio AT2 system.

### Drug treatment

PDEs were derived and immediately injected with approximately 100 ul of triamcinolone (Kenalog-10 triamcinolone acetonide injectable suspension, USP, Bristol Myers Squibb) or vehicle (PBS) using a syringe. PDEs were transferred to 12 well plates and medium was carefully added as described above. After 24 hours in culture, PDEs were collected and RNA extracted as described below.

### RT-qPCR

PDEs were washed with PBS and homogenized in Buffer RLT + BME (1:1000) using the FastPrep-24 Classic bead beating system with lysing Matrix E (MP Bio). The supernatant was transferred to QIAshredder columns and RNA was isolated using RNeasy Mini Kit and RNase-Free DNase set (Qiagen) following the manufactures protocol. cDNA was synthesized using SuperScript II Reverse Transcriptase with Oligo (dT) 12-18mer Primers (Thermo Fisher Scientific) using Mastercycler X50i (Eppendorf) following the manufactures protocols. qPCR was done using Quant Studio 6 Pro (Applied Biosystems) using Fast Start Universal SYBR Green Master Mix (Roche). Primer sequences are listed in Table S1. *ACTB, PPIA,* and *TBP* were evaluated for housekeeping gene stability across drug treatment and patient ID using the normfinder function in the dhammarstrom/generefer (v 0.1.2) package in R (v 4.2.1). *ACTB* was the only gene to meet the stability threshold of < 0.15 for both treatment and patient ID (Table S2) and was used for normalization in subsequent experiments. Relative gene expression was calculated using the ddCt method with *ACTB* used as the housekeeping gene.

### Statistical Methods

All analysis and plotting were done using R (v 4.2.1). For statistical analysis, a linear mixed-effects model with a random intercept for tumor was used to summarize and compare relative gene expression using the lme4 (v 1.1-30) and emmeans (v 1.8.5) packages. P-values were automatically adjusted using the Tukey method. Data were plotted using the ggplot2 (v 3.3.6). For correlation analysis, normality was first assessed by a Shapiro–Wilk normality test. Data following a normal distribution were analyzed by Pearson’s product–moment correlation, while non-normally distributed data were analyzed by Spearman’s rank correlation rho. For the correlation plot, data were fit by the stat_smooth function with method set to lm and plotted using ggplot2. For the correlogram, the rcorr function in Hmisc (v4.7-0) was used to generate the Spearman’s correlation matrix, which was plotted using the ggballoonplot function in the ggpubr (v 0.4.0) package.

## Acknowledgments

We would like to thank all patients who have donated samples for this study. This work was funded by the Children’s Tumor Foundation’s Young Investigator Award (Grant ID: CTF-2021-01-004; https://doi.org/10.48105/pc.gr.146716) and the Van Andel Institute.

---

**Table S1.**
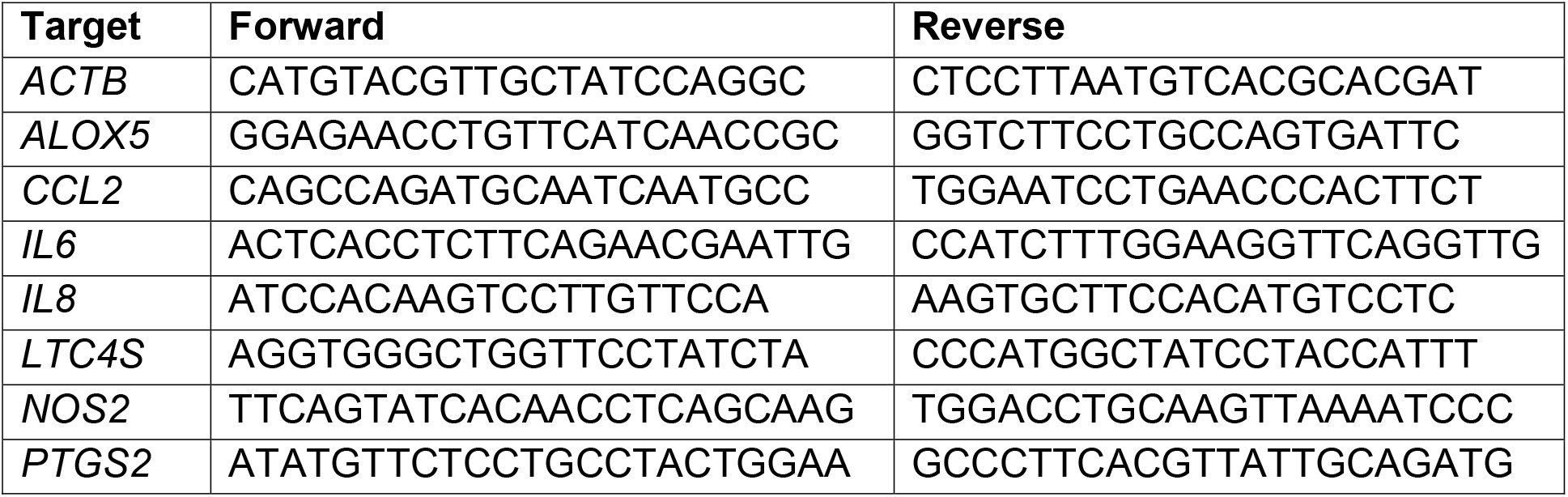
RT-qPCR primers.

**Table S2.** Normfinder housekeeping gene statistics.

| By treatment |  |  |  |
| --- | --- | --- | --- |
| Target | GroupDif | GroupSD | Stability |
| *ACTB* | 0.081 | 0.2 | 0.089 |
| *PPIA* | 0.267 | 0.441 | 0.212 |
| *TBP* | 0.186 | 0.606 | 0.275 |

**Table S2.** Normfinder housekeeping gene statistics.
| By patient ID |  |  |  |
| --- | --- | --- | --- |
| Target | GroupDif | GroupSD | Stability |
| *ACTB* | 0.099 | 0.256 | 0.129 |
| *PPIA* | 0.261 | 0.483 | 0.248 |
| *TBP* | 0.36 | 0.534 | 0.293 |

Supplemental Figure 1. CC3 staining in the epidermis. Representative images (200x) of CC3 positive cells in the stratum basale layer of (a) 3 formalin fixed CNFs and (b) 3 CNF PDEs that were maintained in culture for 1 day.

Supplemental Figure 2. CNF PDEs model inter-tumoral heterogeneity. Representative images of S100B positive tumor cells (a) and CD34 positive dermal fibroblasts (b) in CNF PDEs (20x). Insets (200x) show several regions where S100B and CD34 were inversely expressed, indicating regional tumor heterogeneity of tumor cells and fibroblasts.

