## Supplementary figures and images for "Reciprocal inflammatory signaling in an ex-vivo explant model for neurofibromatosis type 1-related cutaneous neurofibromas"

### Supplemental Figure 1

#### Slide 1
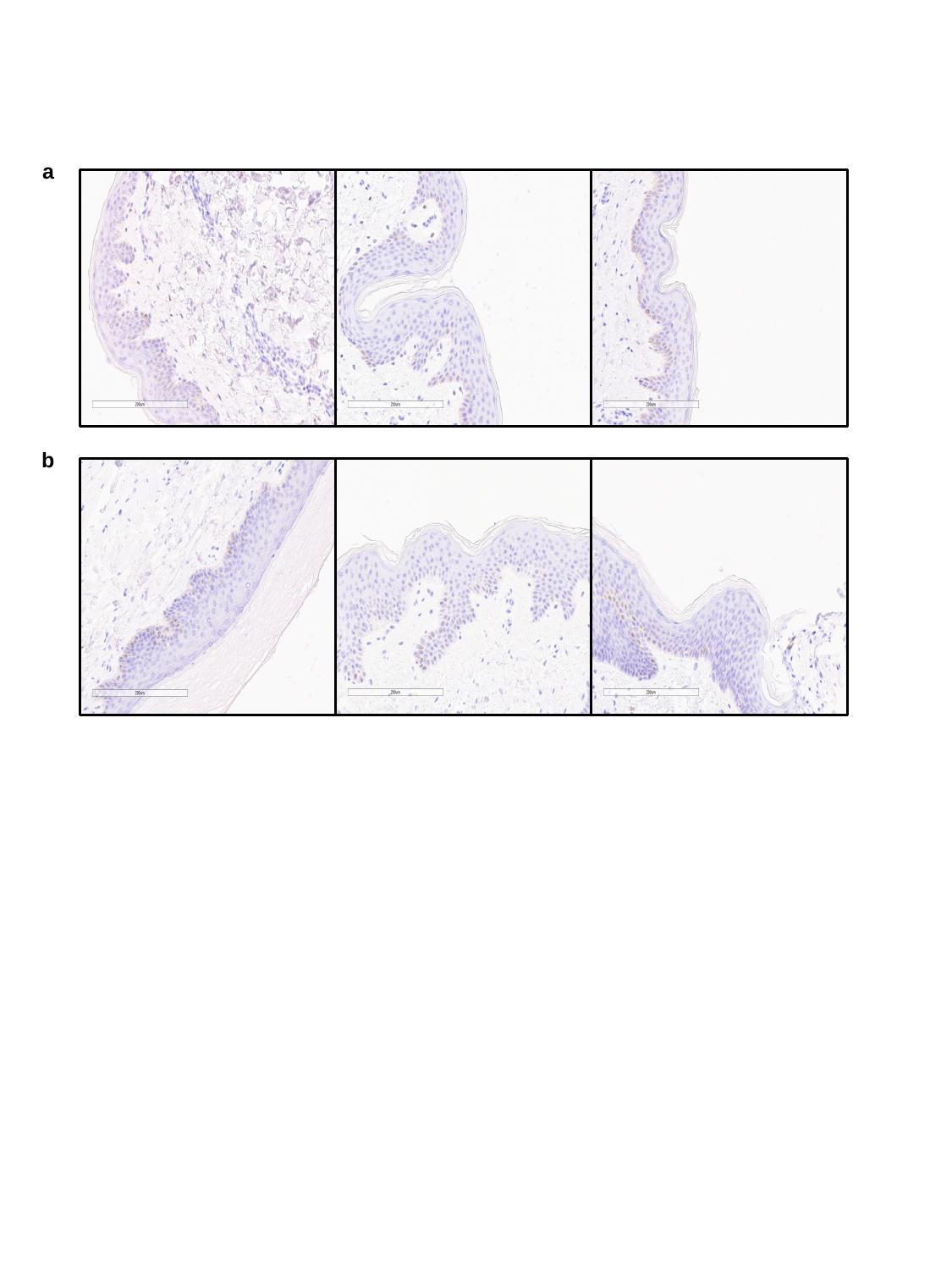

a
b

### Supplemental Figure 2

#### Slide 1
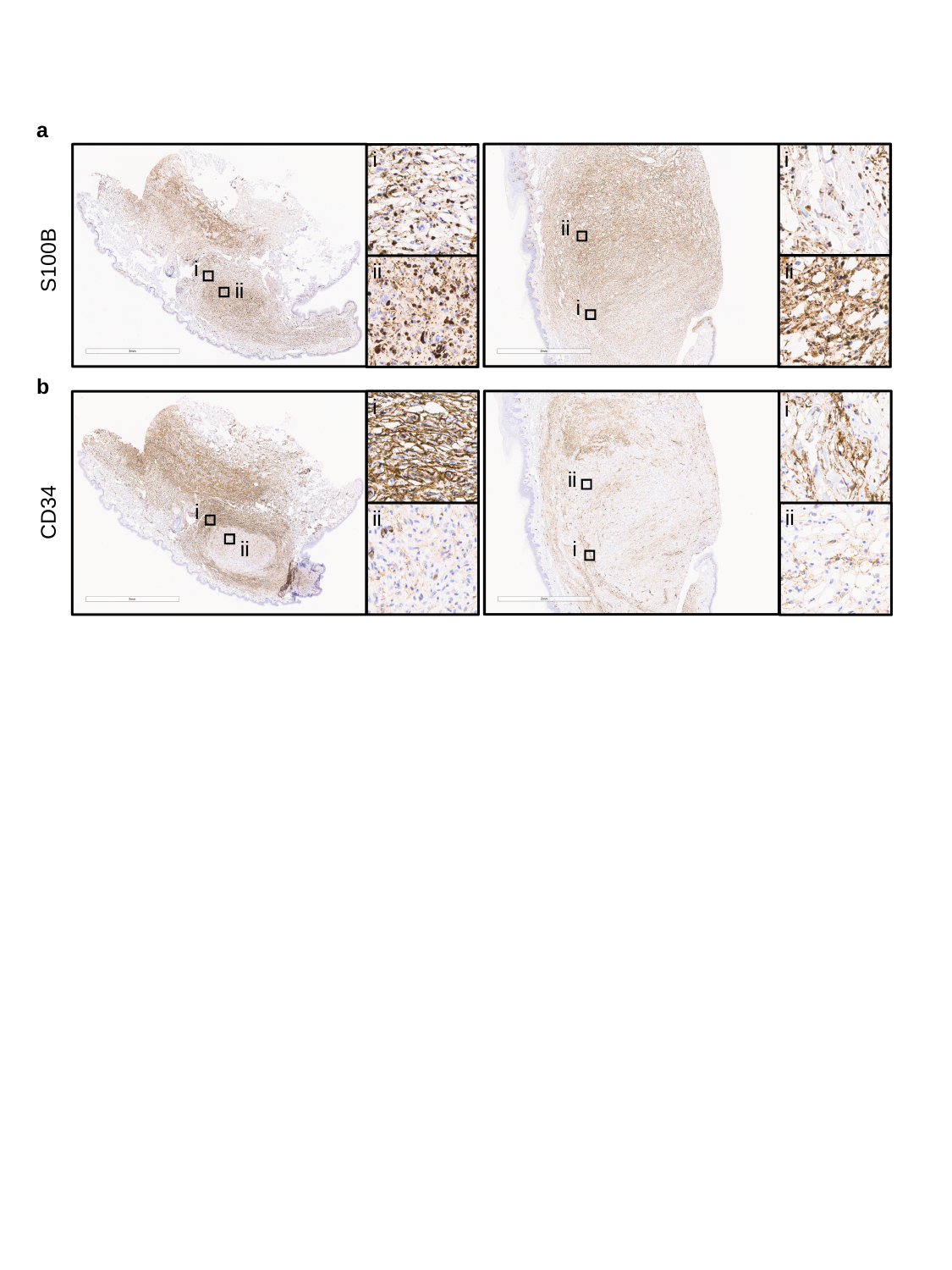

a
i
i
ii
S100B
i
ii
ii
ii
i
b
i
i
ii
CD34
i
ii
ii
i
ii
